# Conservation of non-consensus nucleotides in transcription factor binding sites

**DOI:** 10.64898/2025.12.07.692868

**Authors:** Jenya Belousova, Roberto Morán-Tovar, Malancha Karmakar, Mikhail S. Gelfand

## Abstract

Transcription factors (TFs) affect gene expression by binding to their sites (TFBSs) in the genome. As regulation of transcription is essential for the optimization of metabolism, evolution of TFBSs is thought to be restricted by the TF binding preferences. This means that nucleotides important for binding (consensus nucleotides, CNs) experience the pressure of negative selection and, therefore, are highly conserved. However, for some regulated genes, the strongest possible repression or activation may not necessarily be optimal (i.e. provide the highest fitness). Here, we show that, along with CNs, nucleotides causing relatively weaker binding (non-consensus nucleotides, NCNs) are sometimes preserved by negative selection. We then consider several possible reasons for the conservation of NCNs and demonstrate that NCNs depend epistatically on other loci of the same TFBS for a substantial fraction of TFBSs.

## Introduction

Transcription is a crucial process across the tree of life and is therefore regulated by all organisms at many different levels. Specifically, bacterial RNA-polymerase can interact with a variety of transcription factors (TFs) that bind to specific sites near or intersect with promoters of transcription and activate or inhibit the polymerase’s activity. The TFs’ binding to DNA, in turn, depends on internal or external factors acting on the bacterium at a given moment. Therefore, by adjusting the action of TFs on specific promoters, bacteria can modify gene expression in response to changing conditions or environment.

Bacterial TF binding sites (TFBSs) are short DNA sequences (typically 16-20 bp) usually located upstream of the regulated gene. In a given genome, a TF has a set of similar but not identical TFBSs, and the respective genes form the TF’s *regulon* (Fig. 1(a)). A set of genes regulated by orthologous TFs in related species is called *regulog* (Fig. 1(b)).

**Figure 1.**
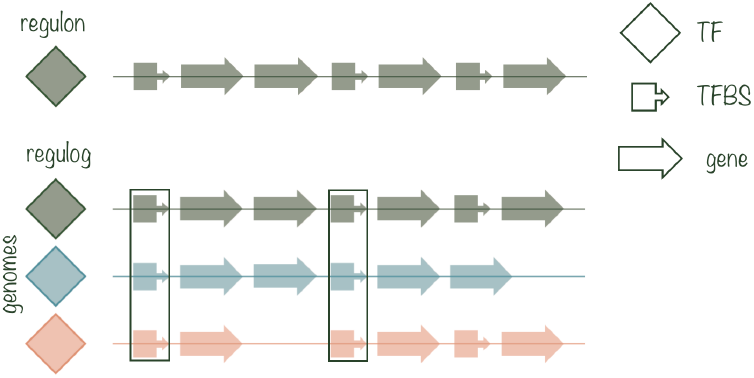
Data structure. Scheme representing TF’s regulon and regulog (bacterial species are shown with different colors). CNs and NCNs are defined based on PWM which is computed over all TFBSs in the regulog. We look at CNs and NCNs conservation within green frames.

TF binding to its TFBS can be characterized by the binding energy *E*_site_. To a good approximation, *E*_site_ is the sum of independent contributions from important positions of the binding site sequence 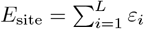, where *L* is the length of the site (Berg et al., 2004). *ε*_*i*_ can be measured experimentally (He et al., 1996; Stormo and Fields, 1998) or calculated given a set of TFBSs (Stormo and Fields, 1998). The importance of a certain nucleotide for binding is reflected in the frequency of this nucleotide across TFBSs of this TF (all sites in Fig. 1(b)). Knowing the frequencies of all nucleotides *N* at each position *i, p*(*N*_*i*_) one can calculate a positional weight matrix (PWM) which can be used to approximate *E*_site_ of binding for a given site sequence *N*_1_…*N*_*L*_:

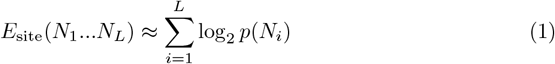

A TFBS position can have one nucleotide whose frequency substantially exceeds the frequencies of other nucleotides. In this case, the prevailing nucleotide is called consensus (CN) one, and three other nucleotides are called nonconsensus (NCN). Some positions do not influence the TF binding, and hence do not have a consensus. The approximation assuming that all CNs contribute equally to *E*_site_ with a value *ε*, while all NCNs have 0 contribution to *E*_site_ is called *the two-state model* (Berg and von Hippel, 1987). CNs can form palindromes and direct repeats, reflecting the symmetrical structure of homodimeric or multimeric TFs (Mazón et al., 2004).

CNs are important for binding and hence are protected from substitutions into NCNs by negative selection. In alignment of a set of orthologous TFBSs (shown with the green frames in Fig. 1(b)), CNs would presumably be conserved. (Note that consensus positions are defined by comparison of TFBSs of different genes in one genome, while here we consider conservation of TFBSs of orthologous genes in different genomes.) However, NCNs can also be conserved (Kotelnikova et al., 2004), even at a level that exceeds the conservation of CNs (Enikeeva et al., 2007). Following the increase in the number of known TFBSs (Novichkov et al., 2013), we consider large regulogs of TFs regulating a variety of metabolic pathways (see Methods, Table 2) in three bacterial families Shewanellaceae, Bacillaceae, and Enterobacteriaceae, with the aim of (1) identifying conserved NCNs; and (2) providing an evolutionary explanation of this conservation.

If an NCN is indeed conserved in orthologous TFBSs, it can be explained in several ways. Our main hypothesis relates to the fitness landscape of the site. The energy of binding between TF and TFBS *E*_site_ provides a quantitative phenotype for the site (Mustonen et al., 2008). The strongest possible binding of a TF upstream of a particular gene is not necessarily the fittest one in a given environment, with some suboptimal binding affinity being biologically more favorable. In terms of the fitness landscape of the TFBS, the case where the strongest binding is preferred corresponds to the *mesa* landscape; while the preferred suboptimal binding corresponds to the *crater* landscape (Berg et al., 2004) (see also the Supplementary Material). Conservation of NCNs might be caused by selection that maintains a certain non-extreme level *E*^∗^ of *E*_site_ as the site evolves.

An alternative possibility is that the TFBS of interest intersects with a regulatory element not yet discovered that includes the conserved NCN; the latter is a conserved CN for this element. This comes from the fact that there are still many undiscovered TFBSs even in the best studied genomes (Trouillon et al., 2023). Another alternative is that, as TFs and TFBSs coevolve (Rodionov et al., 2005), the TF’s binding preferences have changed in a certain clade of the considered TF family. This might mean a change in the pattern of CNs and hence lead to misinterpretation of new conserved CNs as conserved NCNs.

## Results

### Identifying conserved nonconsensus nucleotides

We analyzed TFBSs in three bacterial families: Shewanellaceae, Bacillaceae, and Enterobacteriaceae. We considered 47 Shewanellaceae, 56 Bacillaceae, and 165 Enterobacteriaceae species, and 17, 6, and 14 TFs, respectively. Here we present the result for Shewanellaceae; the results for Bacillaceae and Enterobacteriaceae demonstrate the same trends and are described in detail in the Supplementary Material.

To identify conserved NCNs, one has to compare the NCNs of interest with elements that evolve neutrally. Following the approach of Denisov et al. (2014), we use the third positions of four-fold degenerate codons in the downstream genes as a neutral control. Although these positions are known to be not necessarily neutral (Novoa et al., 2019), in the context of this study the demonstration that NCNs are not under stronger negative selection than synonymous nucleotides is sufficient to claim that NCNs do not evolve neutrally.

To compare the conservation of nucleotides of interest with the control nucleotides, we used two approaches: calculation of the multispecies conservation metric and identification of selection pressure patterns (Denisov et al., 2014).

#### Comparison of multispecies conservation of non-consensus nucleotides with that of synonymous nucleotides

The analyzed phylogeny of the Shewanellaceae family is shown in Fig. 2 (for details see Data preparation in Materials and Methods). For each of 17 TFs we constructed a logo based on the PWM, the FadR logo is shown in Fig. 2 as an example. We used an empiric definition of consensus nucleotides based on nucleotide frequencies and site symmetry: if the frequency of one nucleotide in a given position substantially exceeded the frequencies of others, this nucleotide was considered a consensus one (CN) while three others were considered non-consensus (NCN); at that, the selected CNs should form an (almost) palindromic structure.

**Figure 2.**
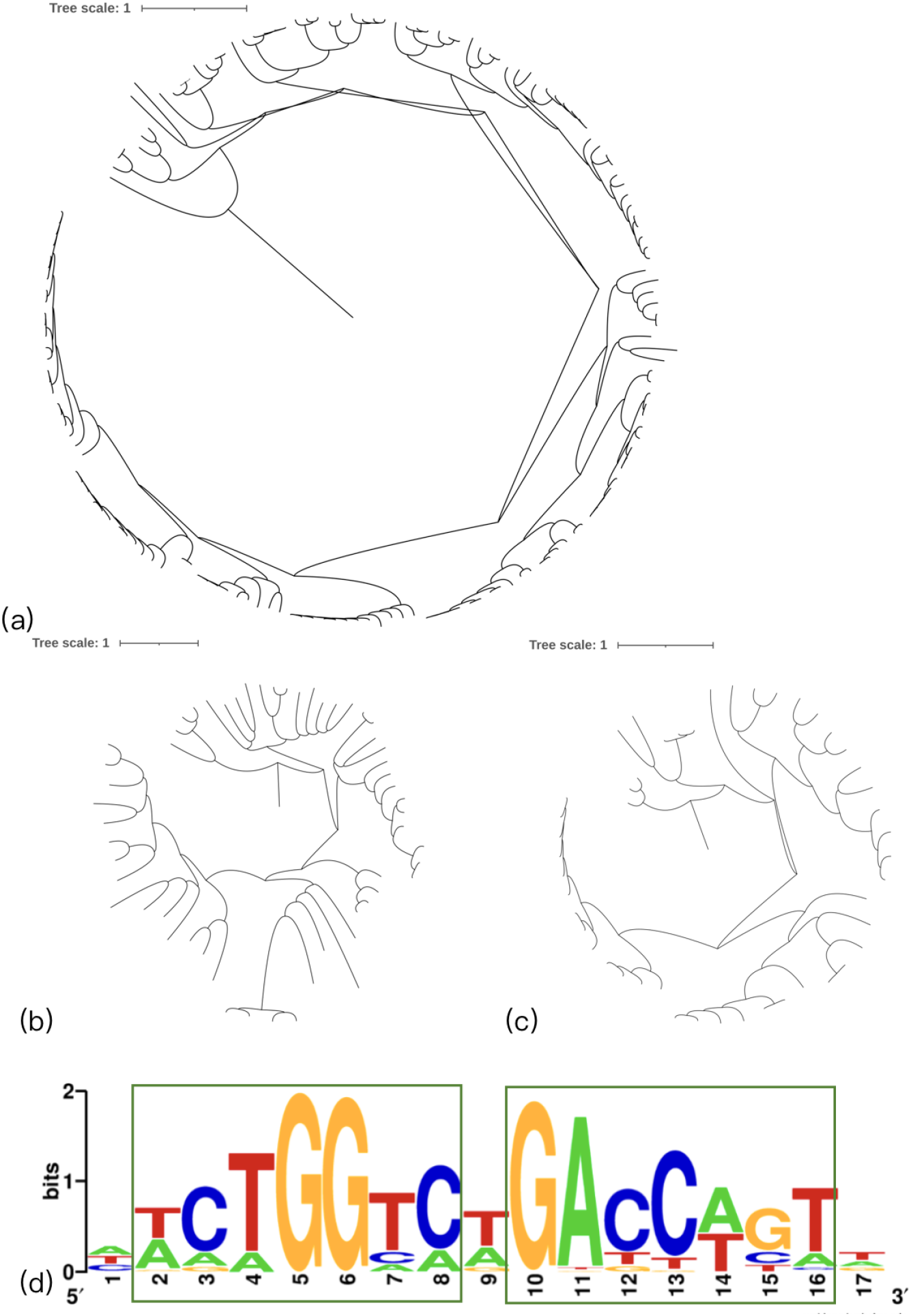
Analyzed bacterial families and the FadR logo. Analyzed phylogenies: **(a)** Enterobacteriaceae; **(b)** Bacillaceae; **(c)** Shewanellaceae. **(d)** Logo (graphical PWM) for FadR in Shewanellaceae. Positions we consider to have a consensus nucleotide are in green frames.

To estimate the rate of evolution of a nucleotide, we calculate the metric of multispecies conservation as follows. For a given alignment with a known phylogeny, we select a reference species *R*. For a given nucleotide in *R* species we measure the phylogenetic distance *D* (as a branch length) between *R* and the most phylogenetically distant species *X* such that this nucleotide is continuously observed in all species *i* such that *D*(*R, i*) ≤ *D*(*R, X*) (Fig. 3a, *D* = the length of the red branch). The conservation of a set of nucleotides can, therefore, be characterized by the distribution of *D* values calculated for all these nucleotides.

**Figure 3.**
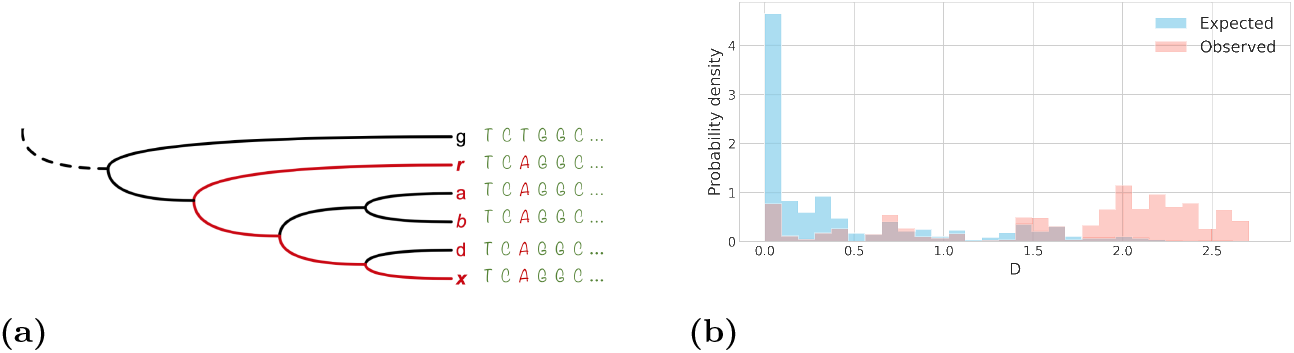
Multispecies conservation metric *D*. **(a)** Calculation of *D* (see the explanation in the text). *D* is equal to the red phylogenetic distance. In this example *D*(*r, a*), *D*(*r, b*), and *D*(*r, d*) are all ≤ *D*(*r, x*). **(b)** Distributions of *D* of NCNs in TFBSs (pink) and neutral controls (blue) for FadR TFBSs in the Shewanellaceae family.

We performed a multi-species conservation analysis for TFs in all three families. The results for Shewanellaceae are shown below; the results for Bacillaceae and Enterobacteriaceae are shown in the Supplementary Material (Fig. 1; Tables 1, 2). In the following, we explain the analysis for FadR in Shewanellaceae as an example, as this factor had the largest site alignments.

**Table 1.** Difference between the distributions of *D* for NCNs and neutral controls for 17 TFs of the Shewanellaceae family.

| TF | U-test | p-value | KL distance |
| --- | --- | --- | --- |
| FadR | 12316691.5 | 0 | 2.03 |
| Crp | 365290419 | 0 | 0.27 |
| ArgR | 2122723 | 6.2e-60 | 0.95 |
| FabR | 2604945 | 4.22e-69 | 0.63 |
| Fnr | 41248189 | 1.02e-269 | 0.72 |
| Fur | 81251121 | 5.74e-157 | 0.23 |
| GcvA | 521590 | 3.55e-22 | 1.24 |
| HexR | 22023578 | 9.2e-240 | 0.77 |
| LexA | 17741337 | 0 | 1.68 |
| MetJ | 6098534 | 1.64e-127 | 0.85 |
| NarP | 8394507 | 5.1e-54 | 0.33 |
| PsrA | 88551141 | 0 | 0.66 |
| TyrR | 6459035 | 1e-198 | 3.47 |
| HypR | 502033 | 2.94e-19 | 0.46 |
| PdhR | 3170190 | 3.85e-51 | 0.73 |
| PrpR | 1667406 | 6.44e-85 | 1.15 |
| SO0072 | 4474417 | 1.85e-74 | 0.58 |

**Table 2.**
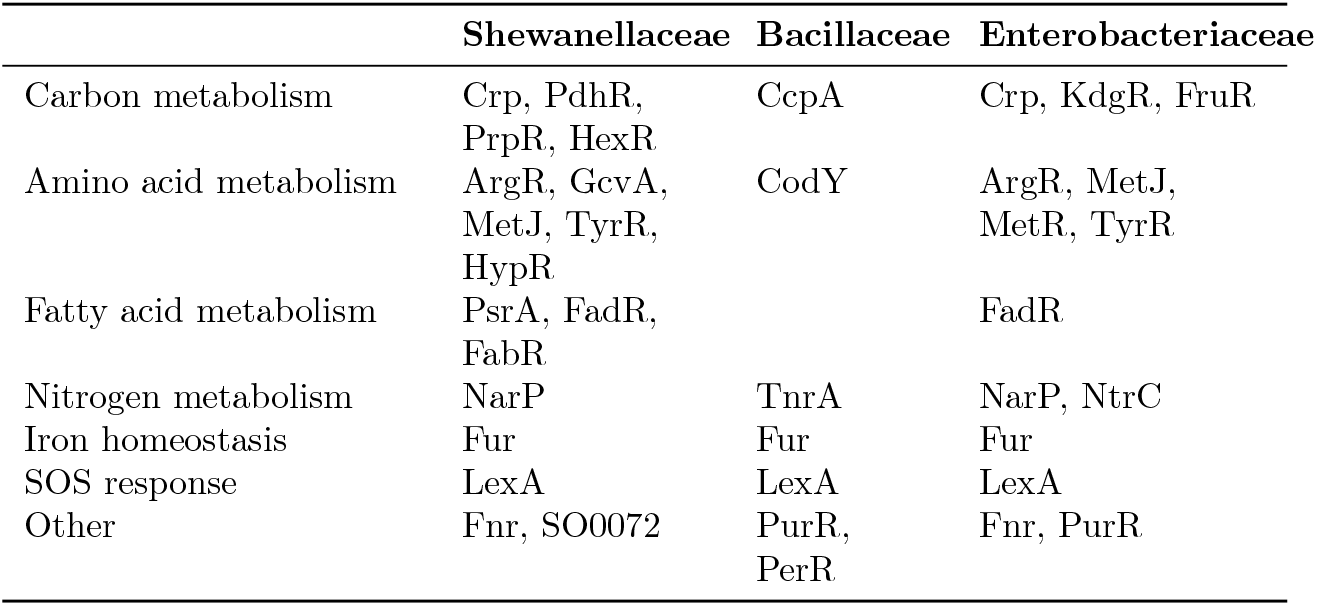
Analyzed TFs.

We considered all alignments of orthologous FadR-binding sites containing ≥ 10 sequences (the sites included in one alignment are schematically shown as vertical frames in Fig. 1). Choosing one genome at a time as reference, we calculated *D* for all NCNs (the pink distribution in Fig. 3b), and for all nucleotides in the corresponding neutral control (the blue distribution in Fig. 3b). The neutral control consisted of all nucleotides in the synonymous codon positions of the downstream FadR-regulated gene, which were the same as observed NCNs in the current reference genome. To compare these two distributions, we used the two-sided Mann-Whitney test and obtained *p* < 10^−324^ (0 in Table 1 refers to *p* < 10^−324^ as well). The *D* distribution for NCNs is therefore significantly shifted towards higher conservation compared to the control distribution, so at least a fraction of NCNs in these TFBSs are indeed more conserved than (almost) neutrally evolving positions. We also calculated the Kullback–Leibler (KL) distance between these distributions (Table 1). Although we do not have reference values for the KL distance to which we could compare our results, the KL distance can be used to compare results for different TFs. For example, the difference between the distributions of *D* for NCNs and neutral controls is especially pronounced for FadR and TyrR. The results for other TFs and families are qualitatively similar (see Supplementary Material).

#### Patterns of selection pressure on CNs and NCNs

Under the basic assumption, consensus nucleotides are crucial for binding of TF to its sites and hence are evolutionary conserved. In terms of the selection coefficient *s*, it means that the consensus nucleotides are protected by negative selection (*s*< 0). On the other hand, NCNs are supposed to evolve under positive selection (*s*> 0) that favors substitutions to CNs. To reveal the pattern of selection acting on CNs and NCNs in our data, we directly measured the value 4*N*_e_*s* (where *N*_e_ is the effective population size) for the substitutions CN → NCN and NCN → CN in all positions (Eq. 5, Methods). The resulting selection pattern for FadR TFBSs is shown in Fig. 4.

**Figure 4.**
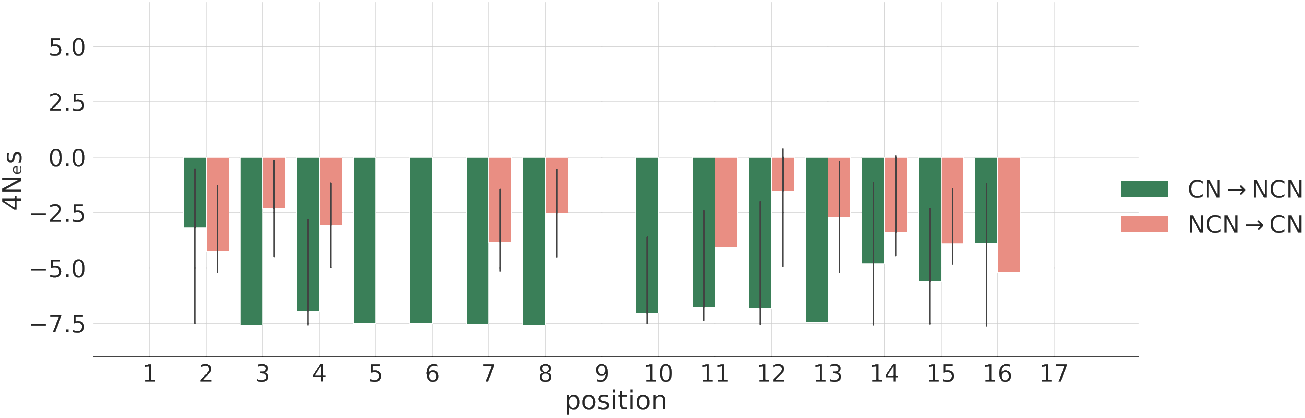
Pattern of selection acting on positions of FadR TFBSs in Shewanellaceae. Selection acting on the CN → NCN and NCN → CN substitutions is shown with green and pink bars, respectively. Two values of 4*N*_e_*s* are calculated for each position in a set of orthologous TFBSs (one vertical frame in Fig. 1). Error bars represent 95% percentile intervals calculated for all sets of orthologous sites regulated by FadR (see Methods).

As expected, negative selection acts on the CNs. However, negative selection of a comparable strength acts on the NCNs as well, meaning that the NCNs are also preserved in evolution.

We repeated this analysis for all other TFs in Shewanellaceae, Bacillaceae, and Enterobacteriacea and observed qualitatively similar patterns of selection (see Supplementary Materials, Fig. 2, 3, 4).

Our results suggest that a substantial fraction of the NCNs are evolutionarily conserved. However, they do not identify the factor(s) that cause this conservation.

### Explaining conservation of nonconsensus nucleotides

We considered potential factors that could cause the conservation of NCNs.

#### Selection acts on the energy of binding between TF and TFBS

The main property of a TFBS that affects its fitness is the energy of binding of TF to it. This means that selection acts directly only on the energy and through this action indirectly constrains the evolution of the site sequence (Mustonen et al., 2008). Our first hypothesis explaining conserved NCNs is that they might be responsible for the conservation of site energy at a certain level larger than the minimum energy (the minimum energy provides the strongest binding). This would allow bacteria to fine-tune the level of TF activity in the transcription of particular genes.

Assume that selection does constrain the changes in the site energy and keeps it at a certain level as the site evolves. This process would narrow the distribution of site energies within a set of orthologous sites (the green frame in Fig. 1). This would also mean that some or all nucleotides are mutually responsible for the conservation of energy, i.e. they interact epistatically. To check whether this is the case, we apply the following procedure. For a set of orthologous sites, we calculate the standard deviation (SD) of their energy distribution. Then, as shown in Fig. 5(a) for a toy alignment of orthologous sites, we shuffle nucleotides within the alignment columns. If NCNs are indeed involved in the conservation of the binding energy, shuffling would destroy putative epistatic interactions between nucleotides within sites. As a result, the standard deviation of the energy distribution would increase. A caveat is that the distribution of site energies might be initially skewed towards large values due to experimental or prediction biases, so we have tested and rejected the possibility of SD growth due to such biases in the data (see Supplementary Materials, Fig. 9).

**Figure 5.**
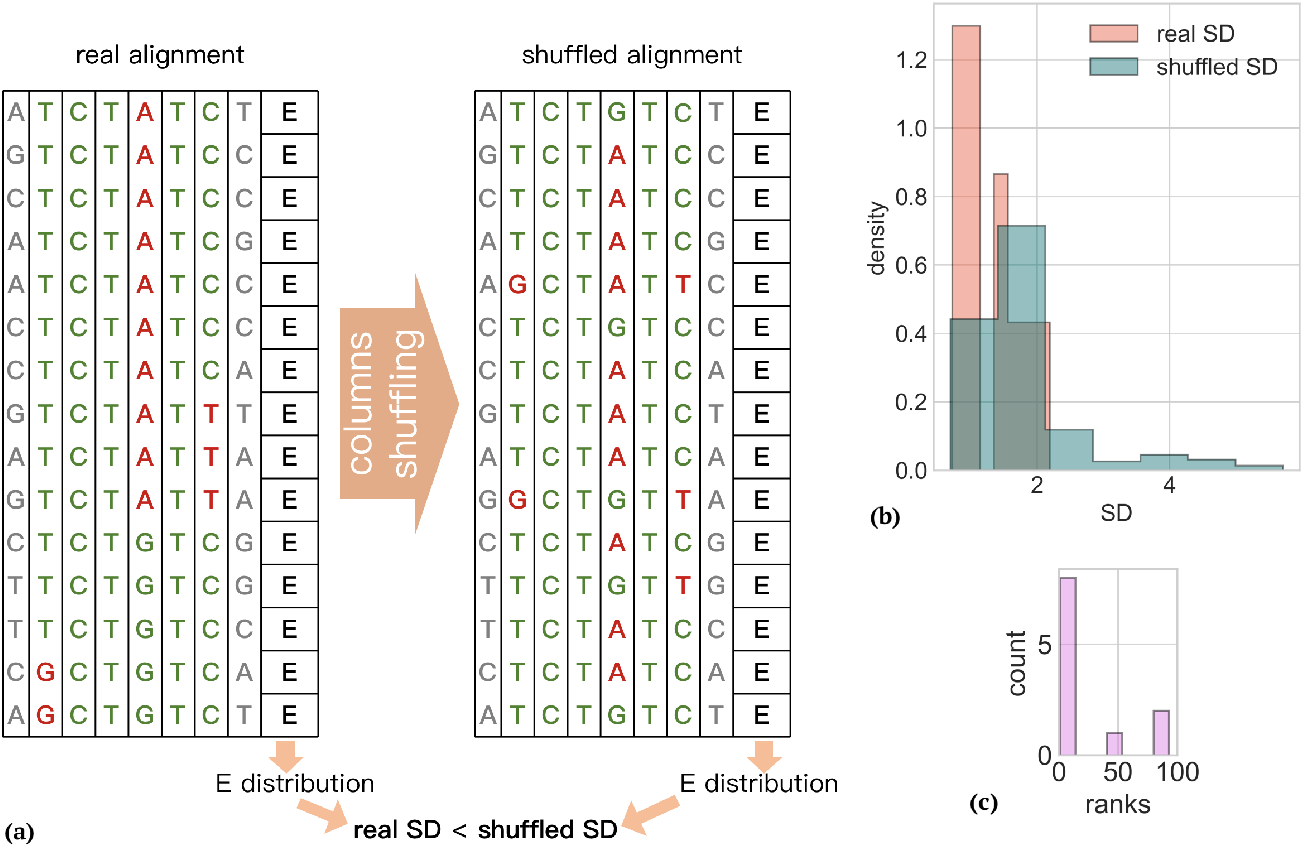
Shuffling procedure for energy distributions. **(a)** Schematic site alignment, where positions without CN are shown in grey, CNs are green, and NCNs are red. After shuffling within alignment columns, the distribution of site energies would become wider if nucleotides within the site interact epistatically; **(b)** Distributions of real SD (pink) and SD after shuffling (blue) for FadR in Shewanellaceae. We used 11 available alignments with 10 and more species; **(c)** Rank distribution computed with SD values represented in (b) (see the explanation in the text).

As an example, we again consider FadR and the alignments of its sites. For each alignment, we calculated the site energy distribution and its standard deviation. The real SDs for 11 alignments of FadR TFBSs form the pink distribution in Fig. 5(b). Then we applied the shuffling procedure to each alignment 100 times, each time calculating SD after shuffling. The resulting distribution of SDs is shown in blue in Fig. 5(b). The two-sided Mann-Whitney test showed significant (*p* = 0.016) difference between these distributions. We also compared these distributions by the rank distribution, where the rank is the position of the real SD value among 100 SD obtained by shuffling, ordered by increase (Fig. 5(c)).

We performed the above procedure for all TFs in three bacterial families. The results of the Mann-Whitney test were not consistently significant for all TFs, and the rank distributions varied. To relate these observations to our previous results, we consider several scenarios of the conservation of NCNs in Supplementary Material (Table 3, Fig. 5, 6, 7, 8).

**Figure 6.**
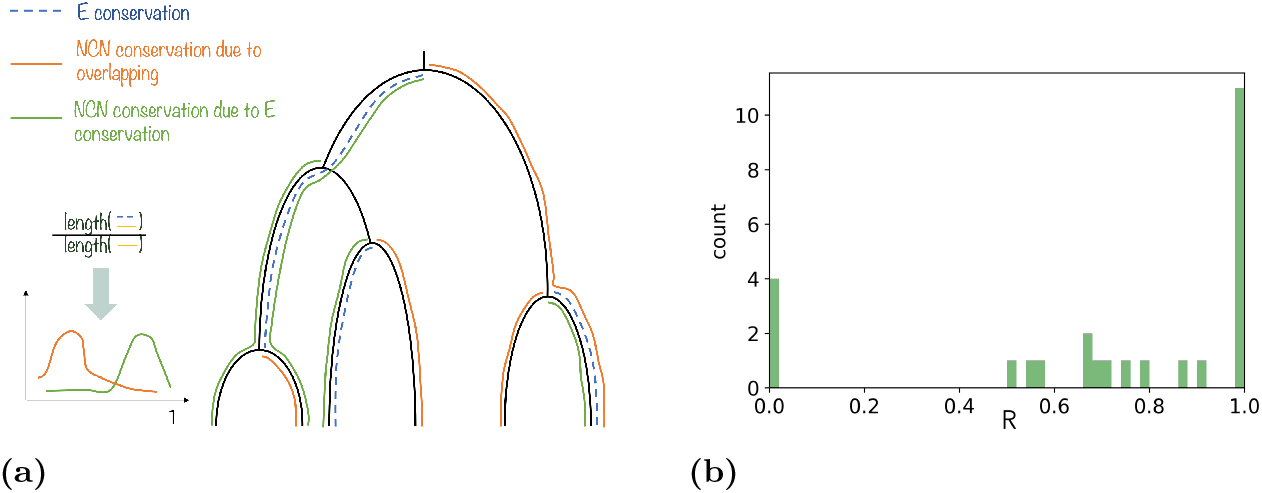
Calculation and distribution of *R*. **(a)** Calculation of the *R* distribution for a hypothetical example. For a given site phylogeny we trace the paths of site energy conservation (blue dashed lines) and paths of conservation for every NCN (solid lines). For every solid line the *R* = *S/B* ratio is calculated to form a distribution. We expect NCNs conserved due to energy conservation to contribute to the distribution in proximity of 1 (the green curve in the hypothetical plot), whereas NCNs conserved due to unknown sites would contribute elsewhere, as they are not linked to the energy conservation (the orange curve); **(b)** The *R* distribution of NCNs for FadR in Shewanellaceae.

**Figure 7.**
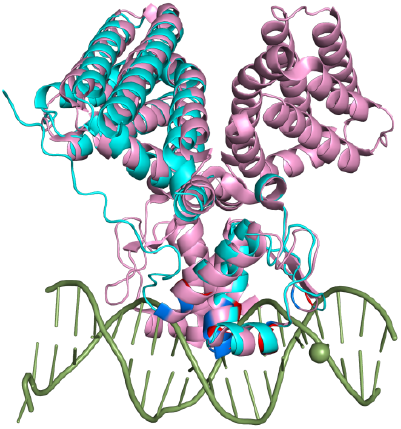
FadR from *Shewanella woodyi* (turquoise) reconstructed by AlphaFold and aligned with FadR from *E. coli* (pink) (1HW2 PDB code). DNA-binding residues of *E. coli* FadR are shown with blue and those of *S. woodyi* FadR are shown with red.

**Figure 8.**
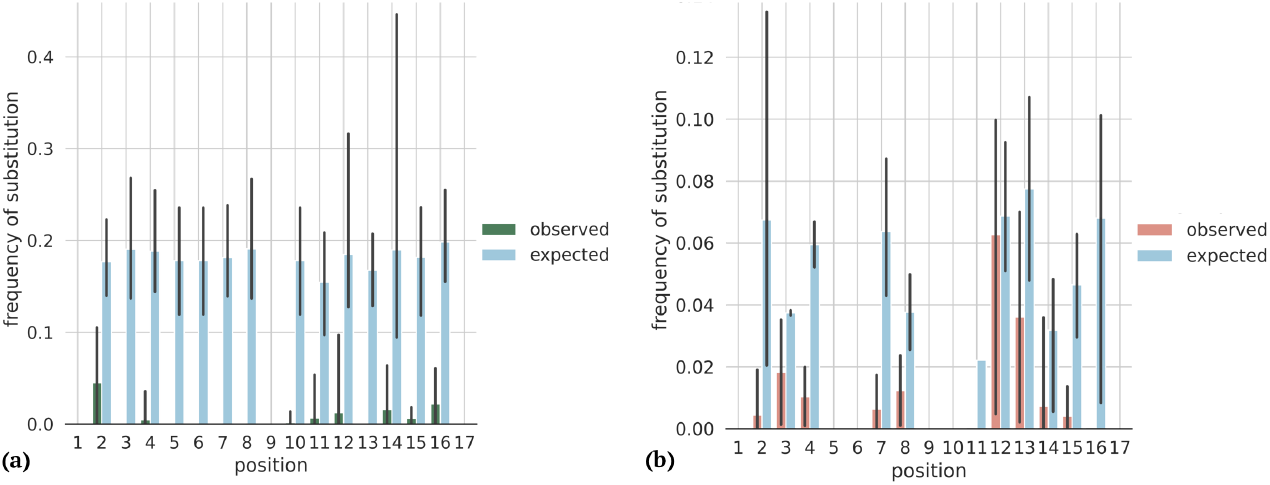
Frequencies of expected (blue bars) and observed substitutions for FadR in Shewanelaceae. **(a)** CN → NCNs, Eq. 3 (green bars); **(b)** NCNs → CN, Eq. 4 (pink bars). Two frequency distributions for every position are calculated within a set of orthologous TFBSs (one vertical frame in Fig. 1). Error bars represent 95% percentile intervals.

#### Overlap with unknown sites

If a site of interest overlaps with an unknown regulatory element (e.g. a site of a different TF), nucleotides that we recognize as non-consensus ones can actually be consensus for these unknown sites. To account for this possibility, we have analyzed how conservation of NCNs is related to conservation of the site energy (Fig. 6a). Given an alignment of TFBSs with known phylogeny and the reconstructed ancestral states, we first traced all phylogenetic paths where the site energy does not change (blue dashed traces). Then, for each NCN we traced the longest path in which this NCN was conserved throughout (solid traces). The evolution of NCNs conserved because of unknown regulatory elements would not be correlated with the energy conservation (orange traces), while the conservation of NCNs responsible for the conservation of energy would match the phylogenetic paths of the energy conservation (green traces). To estimate the dependence of NCN conservation on energy conservation, we calculated the ratio *R* = *S/B* for each conserved NCN, where *B* is the longest phylogenetic distance of NCN conservation and *S* is the overlap of energy conservation with *B*. The distribution obtained for FadR TFBSs is shown in Fig. 6b. As the distribution is shifted towards 1, we conclude that conservation of the majority of NCNs and site energy conservation are indeed correlated, indicating that the NCNs conservation due to unknown regulatory sites is unlikely for FadR. The distributions for other TFs are qualitatively similar in Shewanellaceae, whereas vary between families (see Supplementary Material, Fig. 11).

#### Co-evolution of TFs and TFBSs

TFs evolve, causing changes in TFBS sequences within a certain clade (Korostelev et al., 2016; Suvorova and Gelfand, 2019, 2021). In particular, a change in the TF binding preferences can cause changes in the CN pattern at the site. In this case our approaches would detect conservation of nucleotides assumed to be NCNs which have actually become consensus for a certain clade in the phylogenetic tree. We checked this possibility for FadR. Its crystal structure with defined DNA-binding residues is known for *Escherichia coli* but not for the Shewanellaceae family, so we have reconstructed the 3D structure of FadR from *Shewanella woodyi* using AlphaFold (Jumper et al., 2021) and aligned it to the *E. coli* FadR structure to identify residues interacting with DNA (Fig. 7, the DNA-interacting residues are colored red). Then we aligned the FadR amino acid sequences in the Shewanellaceae phylogeny and observed that all DNA-binding residues were absolutely conserved in all species. This demonstrates that transcription factors do not change their binding preferences significantly within a bacterial family, and our analysis does not misinterpret arising CNs as conserved NCNs.

## Discussion

Our study demonstrates that at least some NCNs are conserved in bacterial TFBSs. Earlier in Denisov et al. (2014), analysis of multispecies conservation *D* demonstrated that NCNs can be conserved at splice sites, although that signal was not detectable by the more direct approach of calculating 4*N*_*e*_*s*. TFBSs demonstrate much stronger conservation of NCNs than expected under neutrality. At least some NCNs evolve under a negative selection of strength comparable to that acting on CNs. This observation is in line with the less direct measurements suggesting such conservation (Kotelnikova et al., 2004; Enikeeva et al., 2007). Negative selection acting on NCNs appears to be ubiquitous, as it has been observed for all studied TFs in three bacterial families. However, the results of two approaches used to find NCNs do not seem to be correlated: TFBSs alignments showing a greater shift in the *D* distributions do not demonstrate stronger negative selection acting on NCNs. The only TF with *D* distribution for NCNs not significantly larger than neutral, PurR in Bacillaceae still demonstrates negative selection acting on NCNs (see Supplementary Material).

We explain conserved NCNs by selection that constrains changes in the binding energy between TF and TFBSs. This was confirmed by shuffling within columns of site alignments. The rank distributions obtained for different TFs always have a substantial fraction of ranks close to 1 which means that at many site alignments the energy *E*_site_ is preserved by selection and kept at some level *E*^∗^. However, the hypotetical scheme in Fig. 5(a) is not the only possible explanation for the widening of the *E*_site_ distribution after shuffling. The increase in SD after shuffling only shows that nucleotides within sites depend on each other, but, in fact, the NCNs responsible for maintaining *E*^∗^ can exist in different positions of orthologous sites, because the same energy level *E*^∗^ can be provided by multiple different site sequences. However, although there are many site sequences that have the same energy *E*^∗^, evolution starting at one of these sequences with particular NCNs to another sequence, with NCNs in different positions, might require a change of *E*_site_ that would considerably deviate from *E*^∗^. Thus, NCNs selected to set a particular *E*_site_ = *E* ∗ might be locked in these positions. This process would depend on how close the actual situation is to the two-state model and on how specific the energy level *E*^∗^ can be. For a substantial fraction of NCNs, we have demonstrated a correlation between their conservation and the conservation of *E*_site_ (Fig. 6) for multiple phylogenies. This observation is in line with the hypothesis that NCNs are responsible for maintaining *E*^∗^ and are therefore conserved at the same site position.

One more approach used to show whether there is a favorable non-minimum level *E*^∗^ for FadR TFBSs is described in the Supplementary Materials. We inferred the dependence of the site fitness on *E*_site_ by scanning the entire genome with PWM, calculating the background energy distribution, and observing deviations from it. One could expect to see a crater fitness landscape (Berg et al., 2004), meaning that the fittest *E*_site_ across the FadR’s regulog is higher than the energy of the strongest binding. However, we observed the mesa landscape (see Supplementary Material, Fig.10) which means that on average sites with minimal *E*_site_ are the fittest. This result is similar to the landscape obtained in Mustonen et al. (2008) for TFBSs in yeast. However, this observation does not disprove that certain genes may favor lower *E*_site_, as this effect might be too subtle to be observed on the fitness landscape calculated for the whole genome—there might be too few true sites to make the dimple detectable.

Our multispecies conservation analysis and the calculation of 4*N*_*e*_*s* demonstrate the existence of conserved NCNs. As it is not possible to directly identify the reason for this conservation, we applied several indirect approaches to estimate the contribution of possible factors to this phenomenon. Firstly, we performed a shuffling procedure and obtained distributions of SD ranks for all TFs. These distributions show that nucleotides within TFBSs depend on each other and often maintain *E*_site_ at a specific level. Secondly, we checked whether the conservation of NCNs is correlated with the conservation of *E*_site_. In several TF families, we observed the conservation of NCNs that was not related to *E*_site_, but for almost all TFs, a substantial fraction of conserved NCNs was associated with conserved *E*_site_. These two observations suggest that NCNs can be responsible for maintaining a fixed *E*_site_ level, which is why they are preserved by negative selection. Lastly, we checked whether our observation resulted from the evolution of TF binding preferences and deduced that TFs were unlikely to change within the considered phylogenies.

Overall, we have demonstrated frequent conservation of nonconsensus positions in binding sites of transcription factors, caused by selection toward optimal (not necessarily minimal) binding energy that causes epistatic interactions between site positions and slows down the evolution.

## Materials and Methods

### Data preparation

Our analysis involved sequences of transcription factor-binding sites (TFBSs) and downstream genes regulated by these sites (Fig. 1). The initial sets of orthologous sites and genes were obtained from the `RegPrecise` database (Novichkov et al., 2013). We considered transciption factors (TFs) with sufficient numbers of TFBSs in genomes from three bacterial families and selected 17 TFs from Shewanellaceae, 7 TFs from Bacillaceae, and 14 TFs from Enterobacteriaceae. These TFs regulate genes involved in a variety of metabolic pathways (Table 2).

The initial sets of orthologous sites and genes were insufficient for the required analysis, so we propagated them as follows. (1) We downloaded complete bacterial genomes from `GenBank` (Sayers et al., 2022). The resulting phylogenies included 47 Shewanellaceae, 56 Bacillaceae, and 165 Enterobacteriaceae genomes. (2) We identified candidate TFBSs with the `z-scan` program (Korostelev et al., 2016) using motifs (positional weight matrices) from `RegPrecise`. The resulting logos for the TFs were obtained using `WebLogo` (Crooks et al., 2004). (3) We resolved the orthology of genes regulated by candidate TFBSs using `ProteinOrtho` (Lechner et al., 2011). As a result, for each TF we obtained sets of orthologous genes and all sites upstream of these genes. The remaining paralogous genes and their sites were excluded from the analysis. Duplicated paralogous sites upstream of the same gene were excluded from the analysis of conservation and selection but included in the calculation of positional weight matrices (PWMs). The PWM value 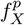 for the nucleotide *X* in position *p* in the alignment of the sites of *m* species was calculated as:

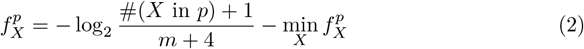

The phylogenies for all three bacterial families were reconstructed using `BV-BRC` (Wattam et al., 2017). As the IDs of the bacterial genomes in `GenBank` and `BV-BRC` did not match, we used the `BV-BRC` search tool to find the genome closest to the one from `GenBank`. The obtained genome sequences matched either precisely or with negligible differences. BV-BRC extracts 100 conserved single-copy genes from Global Protein Families (`PGFams`) (Davis et al., 2016), aligns them, and reconstructs phylogenies based on these genes using `RAxML` (Stamatakis, 2006). For Shewanellaceae, all genes were strictly single-copy in each genome, whereas for Bacillaceae and Enterobacteriaceae, up to ten genomes were allowed to miss a member of a particular homology group and up to ten genomes were allowed to have a duplication of a gene from a homology group. (Gene selection for Bacillaceae and Enterobacteriaceae was less strict so that at least 100 conserved genes could be used.) The trees were rooted by outgroups: *Gallaecimonas xiamenensis* 3-C-1 and *Ruminobacter amylophilus* strain DSM 1361 for Shewanellaceae; *Listeria innocua* ATCC 33091 and *Staphylococcus epidermidis* SK135 for Bacillaceae, *Pasteurella multocida* P2100 and *Vibrio cholerae* N1252 for Enterobacteriaceae. We visualized the trees with `iTOL` (Letunic and Bork, 2024). The trees were processed with the `ETE` toolkit (Huerta-Cepas et al., 2016) in Python.

All alignments were constructed using `MUSCLE` (Edgar, 2004). To align genes regulated by a TF, we aligned the encoded proteins, and then used the induced alignment of synonymous third positions of four-fold degenerate codons as neutral controls. To characterize the patterns of selection given the phylogeny of the species’ phylogeny, we used the baseml program (`model=REV`) and the `codeml` program (`seqtype=codons`, `model=0`) of the `PAML` package (Yang, 2007) to reconstruct the ancestral states of sites and genes, respectively. We observed that `baseml` could reconstruct ancestral states inconsistently given identical inputs. Although this phenomenon had a minor effect on the observations, we attempted to avoid possible artifacts using the following procedure. Firstly, we ran `baseml` for the same site alignment ten times and if all outputs were identical, we considered this output correct. If at least one output was different, we ran `baseml` 100 more times and selected the most frequent output as correct.

### Characterizing patterns of selection pressure on consensus and non-consensus nucleotides

For a set of orthologous sites (one vertical frame in Fig. 1) and regulated genes, we denote the number of direct substitutions from nucleotide *Z* to nucleotide *X* within a given phylogeny by #(*Z* → *X*). Then, for the sites, given all substitutions in every branch of the tree, we calculate the frequency of substitutions from CNs to NCNs at a given position as:

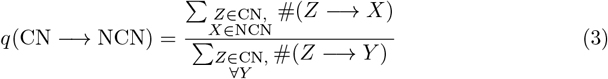

The frequency of NCN → CN substitutions is:

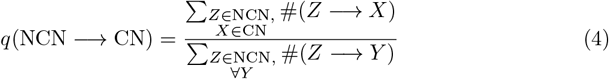

The same calculation was performed for the neutral control, that is, all synonymous four-fold codon positions of regulated gene corresponding to the site. For example, if we encounter a consensus G in a site, we calculate the frequencies of substitutions *q*(G → A | T | C) and *q*(A | T | C → G) for the site (the observed frequency values), as well as *q*(G → A |T| C) and *q*(A |T| C → G) for the neutral control (the expected frequency values).

Thus, for a set of TFBS bound by a given TF, for each site position, we obtain the observed and expected distributions of *q*(NCN → CN) and *q*(CN → NCN). Then we calculate the mean and 95% percentile interval error bars using `sns.catplot()` and `errorbar=(‘pi’, 95)`. This procedure does not require assumptions about the shape of the distribution and shows the spread of *q* (Fig. 8). (The absence of error bars means that *q* was calculated for only one site/gene.)

The probability of fixation of an allele with selection coefficient *s* is 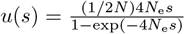, where *N* is the census population size and *N*_e_ is the effective population size (Kimura, 1983). The ratio of substitution rates at loci with the selection coefficient *s* to that at selectively neutral loci with the same mutation rate equals 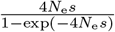, assuming that 4*N*_e_*s* did not change for a long time. These rates were inferred from the data; thus we calculate 4*N*_e_*s* as:

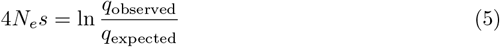

The results for FadR in Shewanellaceae are shown in Fig. 4; the results for other TFs are given in the Supplementary Material (Fig. 2, 3, 4). For each position of a TFBS we divide each value from the distribution of *q*_observed_ (the green distribution in Fig. 8(a) by the mean value of the respective distribution *q*_expected_ (blue in Fig. 8(a) in the same position to obtain 4*N*_e_*s* of the CN → NCN substitutions (the green distribution in Fig. 4). 4*N*_e_*s* of NCN → CN substitutions (pink in Fig. 4) are calculated in the same way using *q*_observed_ and *q*_expected_ from Fig. 8(b). Means and 95% percentile interval error bars in Fig. 4 were calculated analogously using `sns.catplot()` and `errorbar=(‘pi’, 95)` which is a nonparametric approach. All histograms were plotted using the matplotlib.pyplot.hist() function.

## Acknowledgments

This study was supported by the Russian Science Fund, grant 24-14-00276. We thank Professor M. Lässig for insightful discussions that significantly contributed to this work as well as for his guidance and support during this project.

