## Supplementary Material for "Conservation of non-consensus nucleotides in transcription factor binding sites"

### Comparison of multispecies conservation of non-consensus nucleotides with that of synonymous nucleotides

For each TF considered in three bacterial families, we compared the distributions of multispecies conservation $D$, observed for the NCNs and expected for the neutral control. The combined distributions for all TFs in each family are shown in Fig. 1. For Shewanellaceae, the two-sided Mann-Whitney test yielded 2096923012 with $p < 10^{-324}$, and the Kullback–Leibler (KL) distance between these distributions was 0.47. For Bacillaceae, the Mann-Whitney test yielded 52077615 with $p$=$1.94 \times 10^{-81}$, and the KL distance between these distributions was 0.12. For Enterobacteriaceae, the Mann-Whitney test yielded 15426177604 with $p < 10^{-324}$, and the KL distance was 0.49.

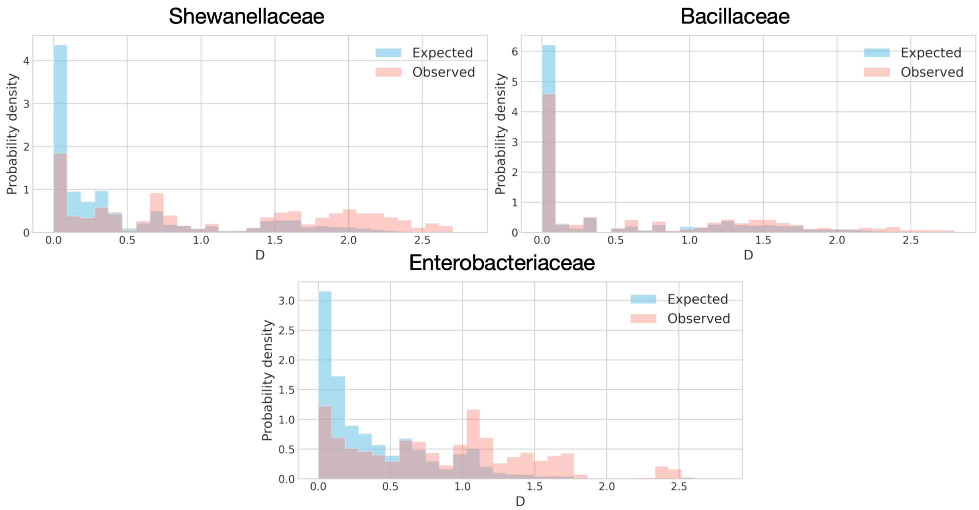


**Figure 1. Distributions of the multispecies conservation metric $D$ for three studied bacterial families.**

For most TFs, the distribution of the multi-species conservation metric $D$ for NCNs shifts towards greater values (Tables 1, 2), although there are exceptions, *e.g.* PurR in Bacillaceae. However, the NCNs of PurR are under negative selection, as can be seen in Fig. 2.

**Table 1. Difference between the distributions of the multispecies conservation metric $D$ for the NCNs and for the neutral control for 7 TFs of Bacillaceae.**

| TF | U-test | p-value | KL distance |
|---|---|---|---|
| Fur | 1020905 | 3.41e-14 | 0.21 |
| PerR | 1156890 | 8.14e-20 | 0.34 |
| PurR | 5366079 | 0.31 | 0.11 |
| LexA | 2002695 | 3.56e-49 | 0.42 |
| TnrA | 10832 | 0.0023 | 3.15 |
| CodY | 435912 | 8.1e-09 | 0.24 |
| CcpA | 3784694 | 2.47e-23 | 0.17 |

**Table 2. Difference between the distributions of the multispecies conservation metric $D$ for the NCNs and for the neutral control for 14 TFs of Enterobacteriaceae.**

| TF | U-test | p-value | KL distance |
|---|---|---|---|
| ArgR | 7406776.5 | 2.59e-71 | 0.66 |
| Crp | 2833714431 | 0 | 0.34 |
| FadR | 20146975 | 0 | 4.13 |
| Fnr | 295148910.5 | 0 | 0.84 |
| FruR | 233854632.5 | 0 | 2.08 |
| Fur | 416648300 | 0 | 0.56 |
| KdgR | 374552259 | 0 | 0.44 |
| LexA | 901056404 | 0 | 0.52 |
| MetJ | 83534704.5 | 0 | 0.7 |
| MetR | 24745448.5 | 1.e-177 | 1.12 |
| NarP | 9948147 | 2.56e-185 | 1.27 |
| NtrC | 27752856.5 | 4.86e-36 | 0.17 |
| PurR | 129884380 | 0 | 0.48 |
| TyrR | 102568550 | 0 | 0.59 |

#### Patterns of selection pressure on CNs and NCNs

The selection that acts on the CNs and the NCNs of all TFBSs is shown in Fig. 2, 3, 4.

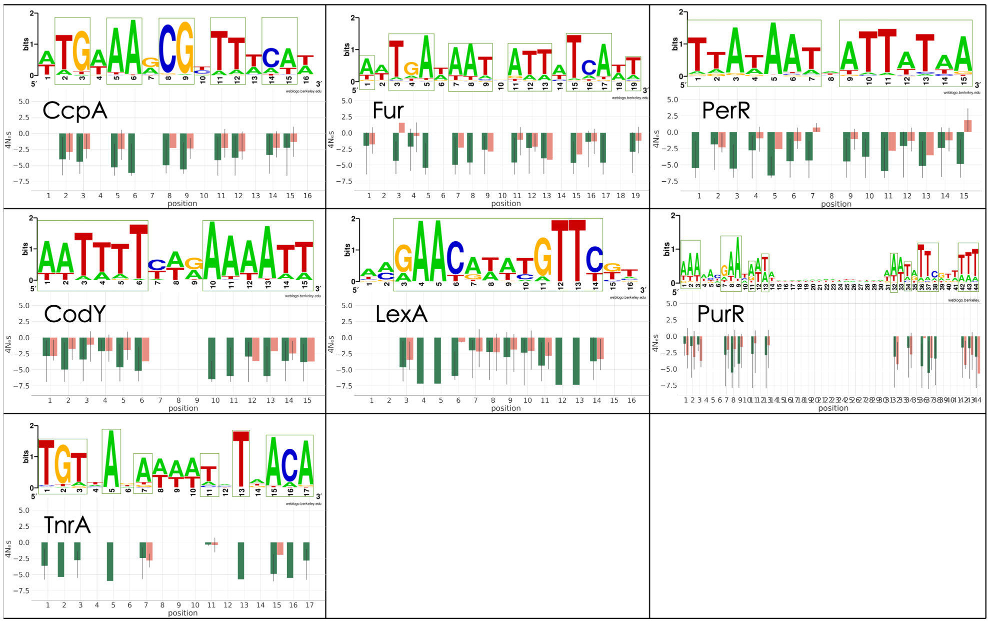


**Figure 2. Logos and patterns of selection acting on TFBSs of studied TFs in Bacillaceae.** Positions containing CNs are in green frames. Notation is same as in the corresponding figure in the main text.

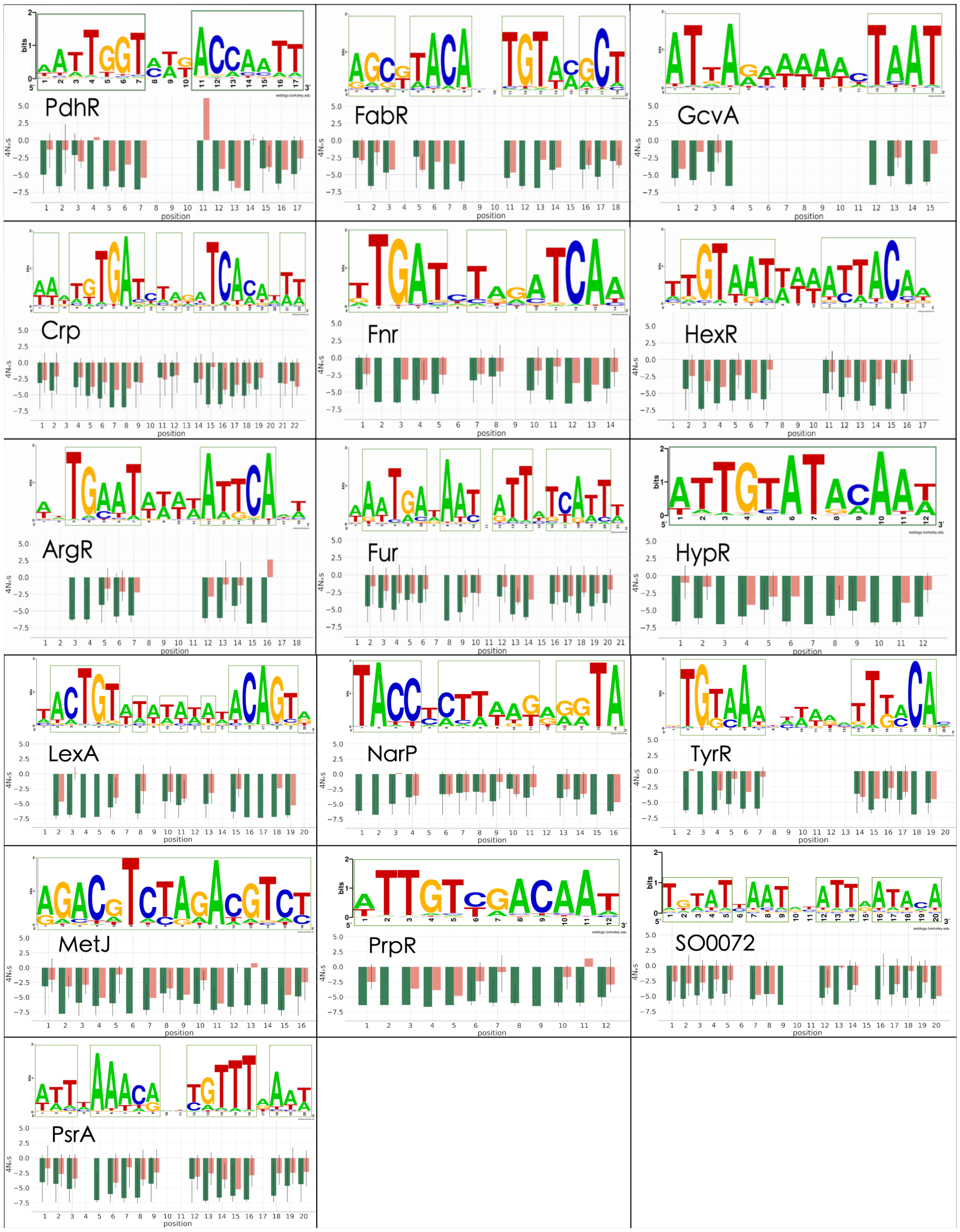


**Figure 3. Logos and patterns of selection acting on TFBSs of studied TFs in Shewanellaceae and logos of the corresponding TFs.** Positions containing CNs are in green frames. Notation is same as in the corresponding figure in the main text.

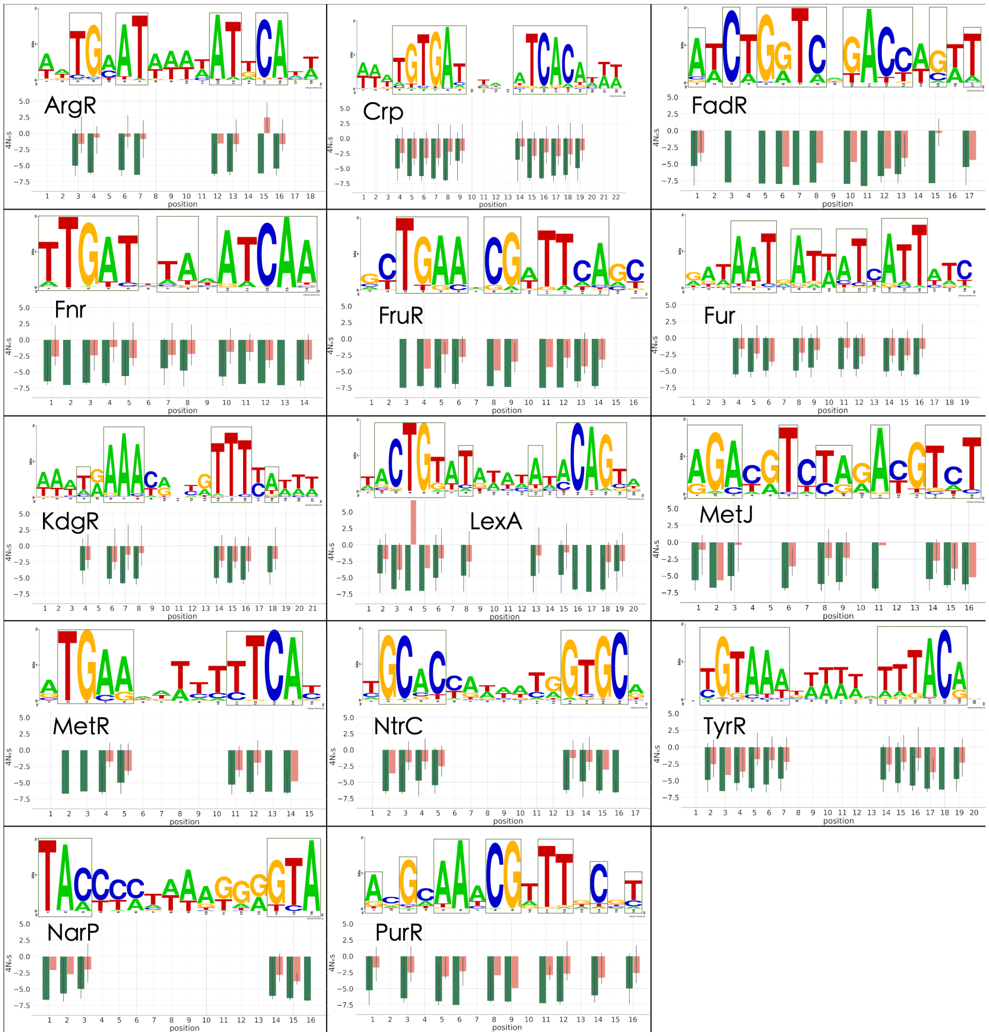


**Figure 4. Logos and patterns of selection acting on TFBSs of studied TFs in Enterobacteriaceae.** Positions containing CNs are in green frames. Notation is same as in the corresponding figure in the main text.

#### Selection acts on the energy of binding between TF and TFBS

We performed the shuffling procedure for all TFs, the results are shown in Table 3, see "Selection acts on the energy of binding between TF and TFBS" in the main text.

**Table 3. Results of the shuffling procedure for all TFs.** U-test value and p-value correspond to two-sided Mann-Whitney test which was applied to rank distributions before and after shuffling procedure, KL distance was also calculated between them.

| TF | U-test | p-value | KL distance | Family |
|---|---|---|---|---|
| HypR | 7586.5 | 0.14 | 0.56 | Shewanellaceae |
| PdhR | 6705 | 0.2 | 0.74 | |
| PrpR | 5478 | 0.57 | 2.78 | |
| SO0072 | 3650.5 | 0.0032 | 1.55 | |
| FadR | 3497 | 0.016 | 0.92 | |
| Crp | 68534.5 | 1.14e-22 | 1.29 | |
| ArgR | 10351 | 0.008 | 2.18 | |
| FabR | 8539 | 0.41 | 0.89 | |
| Fnr | 109823.5 | 2.52e-11 | 0.72 | |
| Fur | 113819 | 3.13e-08 | 0.64 | |
| GcvA | 11374 | 0.13 | 0.3 | |
| HexR | 18124.5 | 0.0017 | 0.9 | |
| LexA | 9660.5 | 0.091 | 1.06 | |
| MetJ | 2118.5 | 0.1 | 2.87 | |
| NarP | 8338 | 0.00039 | 0.94 | |
| PsrA | 58475.5 | 0.0034 | 0.34 | |
| TyrR | 5803 | 0.0085 | 1.29 | |
| CcpA | 4769 | 6.32e-24 | 3.45 | Bacillaceae |
| CodY | 2786 | 0.00025 | 1.15 | |
| Fur | 3068 | 9.51e-06 | 1.04 | |
| LexA | 2260.5 | 0.0028 | 1.26 | |
| PerR | 6193 | 4.93e-05 | 2.3 | |
| PurR | 9567.5 | 0.41 | 1.25 | |
| TnrA | 574.5 | 0.33 | 4.47 | |
| ArgR | 110019 | 5.61e-06 | 0.43 | Enterobacteriaceae |
| Crp | 1490798.5 | 6.74e-31 | 0.42 | |
| FadR | 3480 | 0.47 | 2.25 | |
| Fnr | 376609.5 | 0.00022 | 0.35 | |
| FruR | 48636 | 1.89e-05 | 0.7 | |
| Fur | 544850.5 | 2.58e-12 | 0.59 | |
| KdgR | 805810 | 2.99e-20 | 0.42 | |
| LexA | 400118 | 5.84e-05 | 0.27 | |
| MetJ | 14539.5 | 0.035 | 1 | |
| MetR | 74659.5 | 0.00056 | 0.47 | |
| NarP | 44453 | 7.76e-06 | 0.68 | |
| NtrC | 99978.5 | 2.87e-09 | 0.83 | |
| PurR | 284434 | 0.0059 | 0.17 | |
| TyrR | 68913 | 0.015 | 0.38 | |

Overall, the ranks of real SDs, each corresponding to one set of orthologous sites, fall into one of three categories: close to 1, around 50, or close to 100; the distribution of ranks for each TF is a combination of these three contributions (Fig. 5, 6, 7). To relate this observation to our previous results, we consider several possible cases of the conservation of NCNs, shown in Fig. 8. We assume that only CNs and NCNs make a significant contribution to the change in $E_{\mathrm{site}}$, i.e. that the distribution of $E_{\mathrm{site}}$ is close to the two-state model. Firstly, for some TFs, e.g. PrpR in Shewanellaceae, we observe a prevailing peak around rank = 50, and yet, negative selection is acting on its NCNs (Fig. 3). This is possible if conserved NCNs we observe in several positions have all appeared in different orthologous sites regulated by PrpR. If each site alignment contains at most one conserved NCN, the shuffling procedure would not significantly change the SD of the $E_{\mathrm{site}}$ distribution (because after shuffling there will be the same number of sites with one NCN and the same number of sites without NCNs, as before shuffling, Fig.8(a)). The second possibility arises when NCNs appear in different positions at the roots of non-overlapping subclades, at most one NCN per subclade. Before shuffling, such an alignment would have either none or one NCN per site, but after shuffling, it could have as many NCNs per site as ever emerged in the TFBS tree. As a result, the SD of the $E_{\mathrm{site}}$ distribution would increase after shuffling, and the respective alignment of TFBSs would contribute to the left part of the rank distribution (Fig.8(b)). Finally, the considered phylogeny might be large enough to contain several clades with different optimal levels of $E_{\mathrm{site}}$. For a toy example, if one clade had $x$ conserved NCNs per site, with the rest of the tree having no conserved NCNs, the shuffling procedure would produce NCN counts ranging from 0 to $x$ per site, and the SD of the shuffled alignment would decrease. Such a TFBS phylogeny would contribute to the right peak in the rank distribution (Fig.8(c)).

We studied how these three possible factors affected the rank distributions in the following way. For a given TF, we calculated SDs of the energies for each site alignment and averaged them over the alignment, resulting in the ⟨SD⟩ value. The higher ⟨SD⟩ is, the stronger the first and second factors contribute to the distribution of ranks, and the weaker is the contribution of the third factor. In Fig. 5, Fig. 6, and Fig. 7, we show the rank distributions of all TFs in ascending order of ⟨SD⟩ for the three bacterial families. We can see that the peak around rank 50 is more pronounced for TFs with a lower value of ⟨SD⟩.

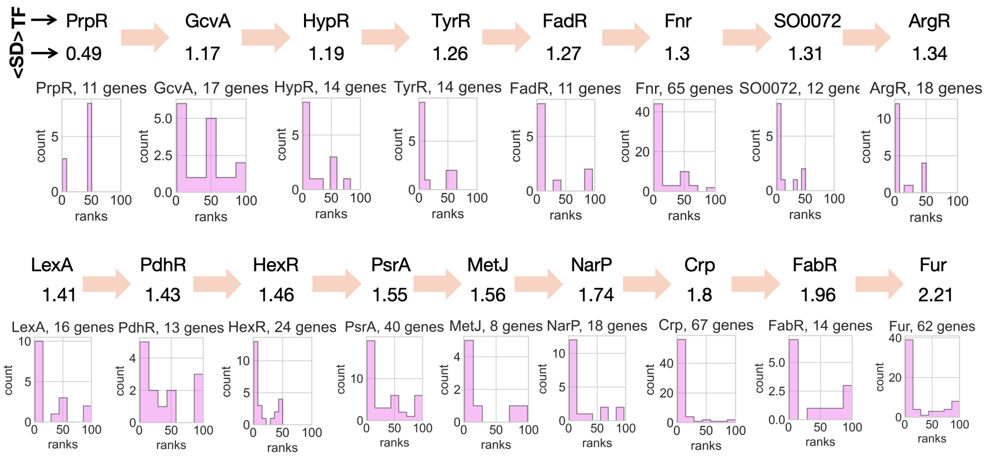


**Figure 5. Distribution of ⟨SD⟩ ranks in the shuffled distribution for all TFs in Shewanellaceae.** (See explanation in the text).

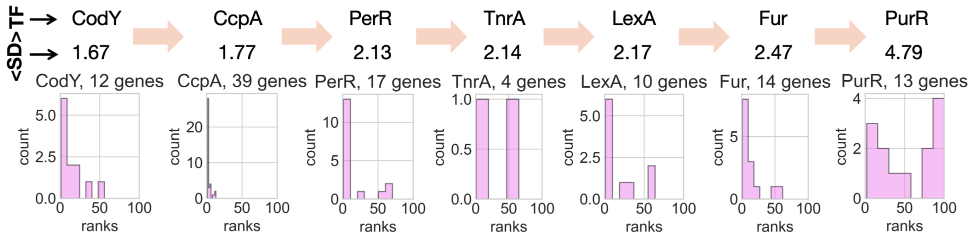


**Figure 6. Distribution of ⟨SD⟩ ranks in the shuffled distribution for all TFs in Bacillaceae.** (See explanation in the text).

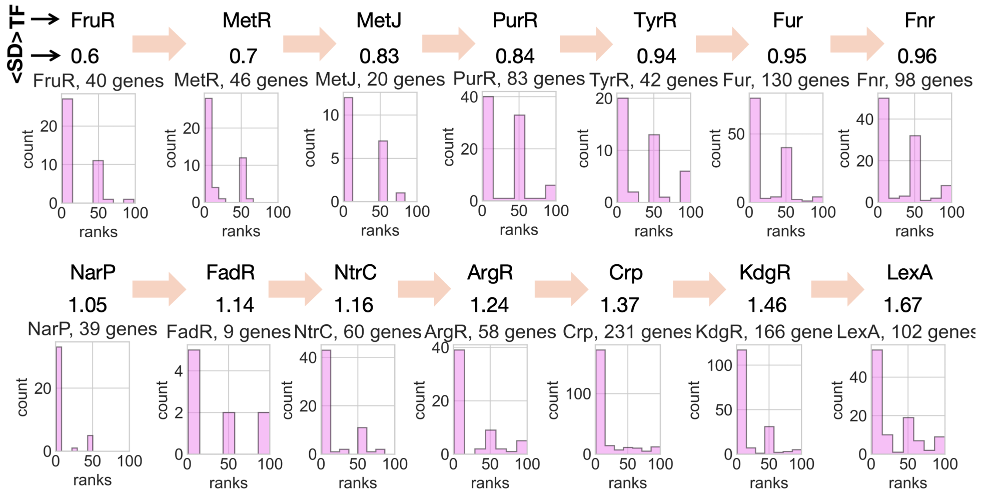


**Figure 7. Distribution of ⟨SD⟩ ranks in the shuffled distribution for all TFs in Enterobacteriaceae.** (See explanation in the text).

| | | | | | | | | | | | | | | | | |
|---|---|---|---|---|---|---|---|---|---|---|---|---|---|---|---|---|
| | T | C | T | G | G | T | C | | G | A | C | C | A | G | T | |
| 1 | 2 | 3 | 4 | 5 | 6 | 7 | 8 | 9 | 10 | 11 | 12 | 13 | 14 | 15 | 16 | 17 |

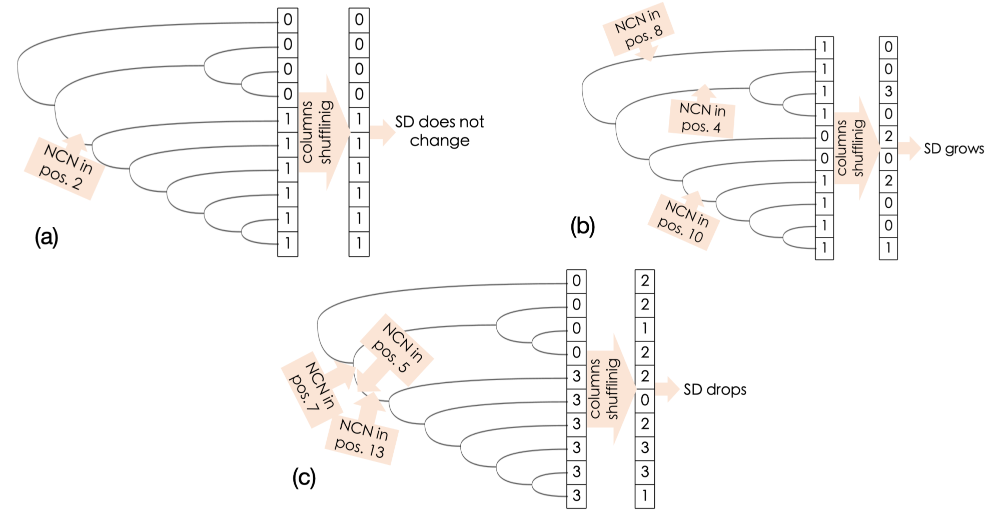


**Figure 8. Possible scenarios of NCNs conservation.** The example TFBS of 17 positions is shown on top with CNs in some positions. Mutations turning CN into NCN occur in this toy phylogeny, and the number at the end of each branch is the number of NCNs in the existing site. The column after shuffling shows hypothetical number of NCNs in the same sites after shuffling. **(a)** Scenario resulting into a peak around 50 for the rank distribution; **(b)** Scenario, resulting into a peak close to 0; **(c)** Scenario, resulting into a peak close to 100.

### Data bias does not affect the results of shuffling

The initial sets of TFBSs in this study come from Novichkov et al. (2013), where the TFBSs were obtained by scanning the genomes with PWMs for the studied TFs. A putative TFBS was accepted if two conditions were met: $E_{\text{site}} >$ threshold, and the site is upstream of a gene orthologous to a gene known to be regulated by the studied TF. We propagated these sets of TFBSs to wider phylogenies by scanning more genomes with PWMs and accepting only sites with $E_{\text{site}} >$ threshold. This means that TFBSs that contribute to the left tail of the $E_{\text{site}}$ distribution are missed. If this effect were substantial, our shuffling procedure would create $E_{\text{site}}$ distributions with relatively larger SD not due to the role of NCNs, but because of this bias in the data. We checked the significance of this bias on the example of FadR. The distribution of $E_{\text{site}}$ for real TFBSs is shown in Fig. 9(a). In Fig. 9(b), (c) we show how the distribution of SD ranks changes if we exclude the left tail of the $E_{\text{site}}$ distribution up to a certain threshold. The distribution of SD ranks, its mean, and its median are robust as we increase the threshold and exclude the tail of the distribution up to $E_{\text{site}} = 8$. Thus, we believe that data bias does not seriously affect our results in the shuffling approach.

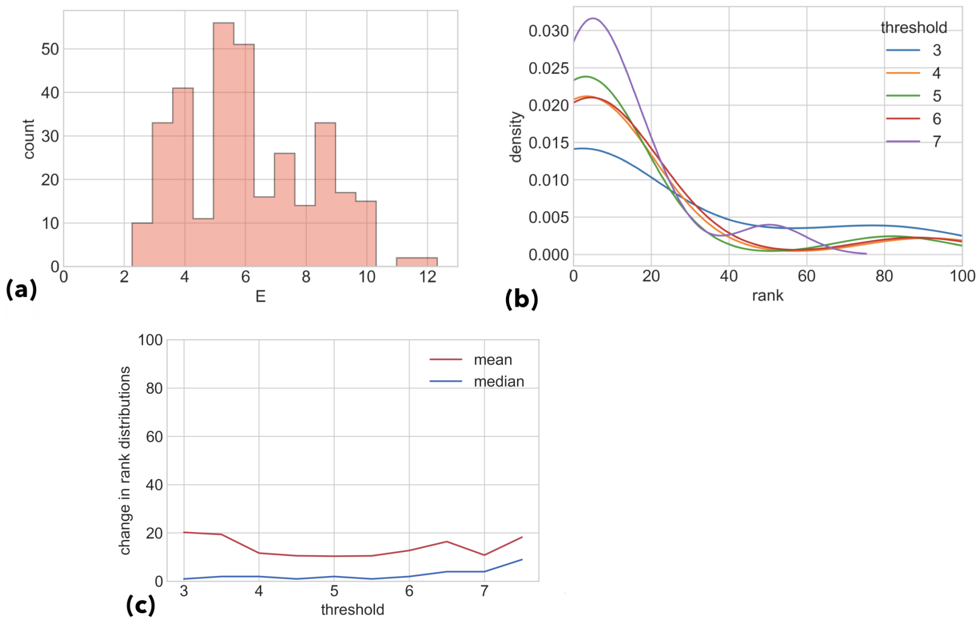


**Figure 9. Effect of data bias on the shuffling procedure for FadR. (a)** Real $E_{\text{site}}$ distribution; **(b)** Change in the distribution of ranks with increasing threshold for $E_{\text{site}}$; **(c), (d)** Corresponding changes of mean and median of the distribution of ranks.

### TFBSs fitness landscape

In the evolution of regulatory sites, energy is more constrained by selection than sequence is. Thus, it is possible to map TFBSs sequences to the phenotype using a quantitative trait, site energy $E_{\text{site}}$ (Mustonen et al., 2008). Consequently, the energy fitness function represents the fitness landscape of the site. We attempted to reconstruct the fitness landscape for TFBSs in the following way (all analyses were performed on FadR in the Shewanellaceae phylogeny). We scanned the intergenic sequences of all genomes with the FadR PWM to construct the histogram of energy counts $Q(E)$ shown in Fig. 10(a).

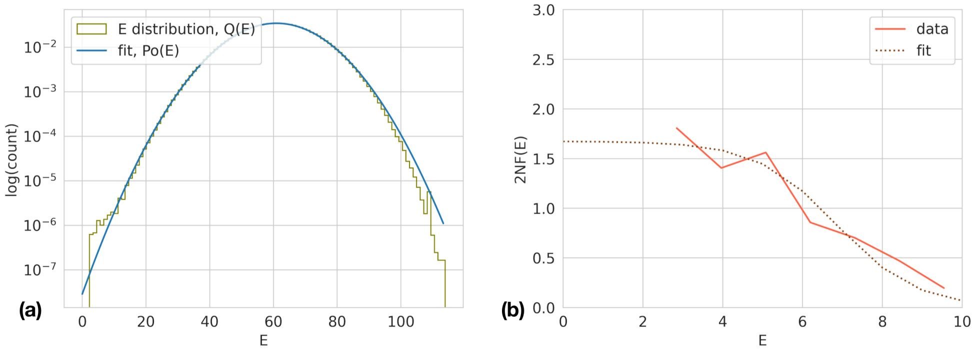


**Figure 10. Inference of the FadR TFBSs fitness landscape. (a)** Histogram of site energies $Q(E)$ predicted by the FadR PWM for all intergenic regions in the Shewanellaceae genomes (olive) and the theoretical Gaussian fit for neutral background sequences $P_0(E)$ (blue). **(b)** Fitness landscape for FadR binding sites inferred from the distributions of $P_0(E)$ and $Q(E)$ (Eq. 1) is shown in pink. The theoretical fit by Eq. 3 is shown with a brown dotted curve.

For $E > 12$ the distribution is close to Gaussian and is well fit by the theoretical Gaussian energy distribution $P_0(E)$, but for $E < 12$ the peak produced by FadR binding sites can be observed. As intergenic sequences are long, using all of them hides this peak, so we used only 800 bp flanks of each intergenic region as genomic background to obtain $P_0(E)$. Varying the length of the flanks does not change the qualitative observations.

Then we inferred the fitness landscape of FadR binding sites. The phenotype distributions $Q(E)$ and $P_0(E)$ are the result of different evolutionary dynamics in the binding sites and in the genomic background, respectively. Neutral evolution leads to the equilibrium distribution $P_0(E)$ of the sequence states, while evolution under selection generates the distribution $Q(E)$. Therefore, the log ratio of these values reproduces the fitness landscape scaled by a factor $2N$ (Berg et al., 2004; Mustonen and Lässig, 2005):

$$2NF(E) = \log \frac{Q(E)}{P_0(E)} + \text{const.} \tag{1}$$

where the constant is arbitrary because only fitness differences matter. The true equilibrium distributions $Q(E)$ and $P_0(E)$ are defined by the changing states of the same locus in time, that is, in the course of evolution, and are not available for observation. Hence, to infer the fitness landscape from Eq. 1, we replace these distributions by the ensemble distributions generated by all functional or background loci for a given transcription factor in a given genome, which we get from our data ($Q(E)$ and $P_0(E)$ in Fig. 10(a)). Thus, we obtain the fitness function $2NF(E)$ shown in Fig. 10(b) (Mustonen et al., 2008).

The naive assumption implies that the minimum binding energy for a site is maintained in the course of evolution and corresponds to the mesa landscape described in Berg et al. (2004) with the sigmoidal function:

$$F(r) = \frac{s}{1 + \exp\left[\varepsilon(r - \rho)\right]} \tag{2}$$

where $s$ is the selection coefficient, $\varepsilon$ is the binding energy per nucleotide mismatch, $\varepsilon\rho$ is the chemical potential measuring the TF concentration, and $r$ is the Hamming distance representing the number of mismatches. However, if the minimal energy is not the fittest, the crater landscape will be observed:

$$F(r) = \frac{s}{1 + \exp\left[\varepsilon(r - \rho_{\mathrm{ON}})\right]} - \frac{s}{1 + \exp\left[\varepsilon(r - \rho_{\mathrm{OFF}})\right]} \tag{3}$$

where the ON and OFF states, respectively, activate and suppress gene expression.

In the FadR example, we see the mesa landscape, suggesting that there is no suboptimal binding level, which would be fitter than the strongest binding for FadR TFBS. Although we have observed conserved NCNs and their correlation with the conservation of $E$, the reconstruction of the fitness landscape of TFBSs does not capture this effect.

### Overlaps with unknown sites

To estimate the dependence of NCN conservation on energy conservation, we calculated the ratio $R = S/B$ for each conserved NCN, where $B$ is the longest phylogenetic distance of NCN conservation and $S$ is the overlap of energy conservation with $B$ (see the main text, "Overlaps with unknown sites"). We obtained the distributions of the ratio $R$ for all studied TFs, shown in Fig. 11. In all three families, we observed a fraction of NCNs whose conservation was correlated with the energy conservation. In Shewanellaceae and Enterobacteriaceae, this fraction was the most significant among all NCNs; however, in Bacillaceae, apart from this fraction, we also observed a large fraction of NCNs which are not correlated with energy conservation.

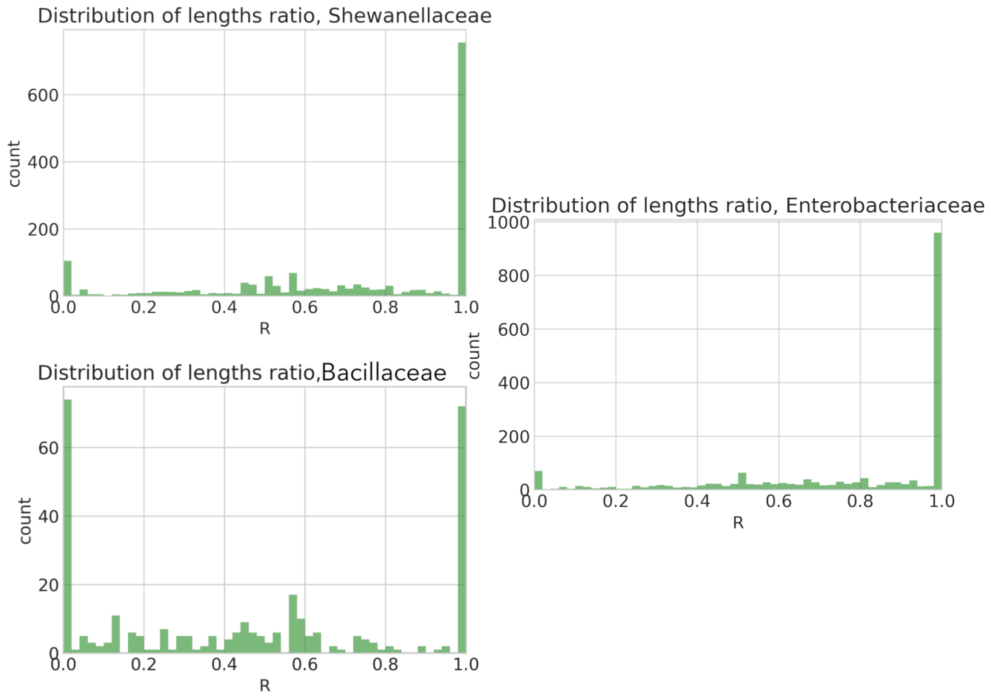


**Figure 11. Distribution of the ratio $R$ for three studied bacterial families.**
